# Colorectal cancer mutational profiles correlate with defined microbial communities in the tumor microenvironment

**DOI:** 10.1101/090795

**Authors:** Michael B. Burns, Emmanuel Montassier, Juan Abrahante, Sambhawa Priya, David E. Niccum, Alexander Khoruts, Timothy K. Starr, Dan Knights, Ran Blekhman

**Author notes:** Corresponding authors (MBB) (RB). Current affiliation: Department of Biology, Loyola University Chicago, Chicago, IL, 60626; USA.

## Abstract

Variation in the gut microbiome has been linked to colorectal cancer (CRC), as well as to host genetic variation. However, we do not know whether, in addition to baseline host genetics, somatic mutational profiles in CRC tumors interact with the surrounding tumor microbiome, and if so, whether these changes can be used to understand microbe-host interactions with potential functional biological relevance. Here, we characterized the association between CRC microbial communities and tumor mutations using microbiome profiling and whole-exome sequencing in 44 pairs of tumors and matched normal tissues. We found statistically significant associations between loss-of-function mutations in tumor genes and shifts in the abundances of specific sets of bacterial taxa, suggestive of potential functional interaction. This correlation allows us to statistically predict interactions between loss-of-function tumor mutations in cancer-related genes and pathways, including MAPK and Wnt signaling, solely based on the composition of the microbiome. These results can serve as a starting point for fine-grained exploration of the functional interactions between discrete alterations in tumor DNA and proximal microbial communities in CRC. In addition, these findings can lead to the development of improved microbiome-based CRC screening methods, as well as individualized microbiota-targeting therapies.

**Author summary:** Although the gut microbiome - the collection of microorganisms that inhabit our gastrointestinal tract - has been implicated in colorectal cancer, colorectal tumors are caused by genetic mutations in host DNA. Here, we explored whether various mutations in colorectal tumors are correlated with specific changes in the bacterial communities that live in and on these tumors. We find that the genes and biological pathways that are mutated in tumors are correlated with variation in the composition of the microbiome. In fact, these changes in the microbiome are consistent enough that we can use them to statistically predict tumor mutations solely based on the microbiome. Our results may be used to understand the roles of specific microbes in CRC biology, and could also be the starting point of microbiome-based diagnostics for not only detection of CRC, but characterization of tumor mutational profiles.

## Introduction

The human gut is host to approximately a thousand different microbial species consisting of both commensal and potentially pathogenic members[1]. In the context of colorectal cancer (CRC), it is clear that bacteria in the microbiome play a role in human cell signaling[2–11]; for example, in the case of CRC tumors that are host to the bacterium *Fusobacterium nucleatum*, the microbial genome encodes a virulence factor, *FadA*, that can activate the β-catenin pathway[12]. In addition, several attempts have been made to predict CRC status using the microbiome as a biomarker[13–16]. It has been shown that by focusing on *F. nucleatum*, it is possible to predict some clinically relevant features of the tumor present[17]. There is a positive, but not statistically significant, association between *F. nucleatum* presence in colorectal cancers in patients who eat a Western diet[18]. Regardless, as only a minority of CRCs are host to *F. nucleatum*, the use of this species as a sole maker are limited[18,19]. Other specific microbes have been linked to CRC, including *Escherichia coli* harboring polyketide synthase (pks) islands[20,21] and enterotoxigenic *Bacteroides Fragilis* (ETBF)[22–24]. The mechanism of action of these associations is still under investigation with *F. nucleatum* being the most clearly developed[12].

A major challenge in the research on the CRC-associated microbiome is the variability of the findings by different research groups. Comparisons among and between the findings from groups studying different patient populations arrive at differing sets of microbes that appear correlated with CRC. As highlighted above, several groups have focused on individual microbial taxa to identify functional associations that explain the statistical correlations between individual microbial species and cancer. It is clear that not all of the structural and compositional changes in the tumor-associated microbiome are functionally relevant and clinically actionable. This situation is analogous to passenger and driver mutations in cancer. In this case there are passenger microbes that show altered abundance at the site of the tumor simply due to the changes brought about by the aberrant physiology of tumorigenesis. Others, the driver microbes, are potentially oncogenic, drive tumor development once it has formed, or some combination of the two. Conversely, in the cases where specific microbial taxa are depleted at the site of the tumor, the microbes may have anticancer effects that remain to be explored.

We know that in healthy individuals, host genetic variation can affect the composition of the microbiome[25–30], and the associated human genetic variants are enriched with cancer-related genes and pathways[26]. However, it is still unknown whether somatic mutations causing disruptions in genes and pathways in the host’s cells can affect the composition of the microbiome that directly interacts with these tissues. It is also clear that individual taxa are likely to have differential interactions with host tissues dependent on the larger context of the community as a whole (*e.g.* not all patients with high levels of *F. nucleatum* in their gut microbiomes will develop CRC). It is likely that the genetic and phenotypic heterogeneity of tumors also results in differential interactions with the microbiome. This variation, if not accounted for when assessing the tumor microbiome, might allow researchers to uncover generic or widely prevalent microbial changes at the site of the tumor. Including genetic information about the cancer in the assessment of tumor-microbiota interactions would allow for a more fine-grained analysis that unmasks the subtle interactions that might be lost in a generic tumor-normal comparison. To that end, we have performed a inter-tumor analysis accounting for the genetic heterogeneity present between them. In this work we show (i) associations between the tumor microbiome and variation in somatic mutational profiles in CRC tumors; (ii) which host genes and bacterial taxa drive the association; (iii) how these patterns can shed light on the molecular mechanisms controlling host-microbiome interaction in the tumor microenvironment; and (iv) how this correlation can be used to construct a microbiome-based statistical predictor of genes and pathways mutated CRC tumors. These findings provide a framework for discovery of sets of microbial taxa (communities) that should be further explored using direct functional assessment to determine passenger or driver status.

## Results

### Changes in the microbiome reflect tumor stage

We performed whole-exome sequencing on a set of 88 samples, comprised of 44 pairs of tumor (adenocarcinomas) and normal colon tissue sample from the same patient, with previously characterized tissue-associated microbiomes[2]. The mutations in each of the tumors’ protein-coding regions were identified relative to the patient-matched normal sample and annotated as either synonymous, non-synonymous, or loss-of-function (LoF) mutations (S1 Table, S2 Table, S1 Fig, and S2 Fig). The ranges of mutations per tumor in our data set were between 19-6678 (total), 10-3646 (missense) and 0-208 (LoF).The mutations were collapsed by gene as well as by pathways using both Kyoto Encyclopedia of Genes and Genomes (KEGG) and pathway interaction database (PID) annotations[31–34] (see Materials and Methods). We performed quality control of the data and stringent filtering at every step, with the goal of reducing false positives in mutation calling and statistical prediction (*e.g.*, requiring 30x coverage at a site in both the tumor and matched normal sample to call a mutation; see Materials and Methods). While these requirements are likely to increase the frequency of false negatives (true mutations that simply do not meet our criteria), this rigorous strategy is appropriate as a means of increasing the biological relevance of our findings. Of note, when comparing the common LoF mutations found in our dataset to those found in colorectal tumors sampled as part of The Cancer Genome Atlas (TCGA) project, we find several commonalities, including a high frequency of LoF mutations in *APC,* as well as numerous missense mutations in *KRAS, NRAS,* and *TP53*, as expected (S1 Table)[35]. In general, the range of mutations across our sample set were also in line with those identified as part of TCGA and other CRC exome sequencing studies (see S15 Table and S16 Table for comparisons)[35–39].

We first investigated the relationship between microbial communities and tumor stage (**Fig. 1**). We hypothesize that the structure and composition of the associated microbiome can be affected by relevant physiological and anatomical differences between the tumors at different stages that would provide different microenvironmental niches for microbes. We identified the changes in the microbial communities surrounding each tumor as a function of stage by grouping the tumors into low stage (stages 1-2) and high stage (stages 3-4) classes, due to the low number of total tumors with available stage information, and applied linear discriminant analysis (LDA) effect size (LEfSe) to the raw operational taxonomic unit (OTU) tables corresponding to these tumors (S3 Table, S5 Table, and S6 Table)[40]. The set of taxon abundances was transformed to generate a single value representing a risk index classifier for membership in the low-stage or high-stage group (**Fig. 1A**; see Methods). To ascertain the error associated with these risk indices, a leave-one-out (LOO) cross-validation approach was applied. We also used the LOO results to generate receiver operating characteristic (ROC) curves and to calculate the area under the curve (AUC; see **Fig. 1B**). In addition, we performed a permutation test to assess the method’s robustness (S5 Table). Using this approach, we demonstrate that the changes in abundances of 31 microbial taxa can be used to generate a classifier that distinguishes between low-stage and high-stage tumors at a fixed specificity of 80% and an accuracy of 77.5% (P =0.02 by Mann-Whitney U test, and P = 0.007 by a permutation test; S5 Table). The resulting changes seen in our analysis of the microbial communities that vary by tumor stage were similar to those found in previous studies, including one using a Chinese patient cohort[4,41]. In both cases, there were significant changes among several taxa within the phylum *Bacteroidetes*, including *Porphyromonadaceae*, and *Cyclobacteriaceae* (**Fig. 1** and S5 Table). We applied the model generated from our data to two independent datasets that were analyses of the CRC microbiome and that also reported tumor staging[10,42] (S3 Fig). In the case of Flemer *et al.* the same approach successfully separated the low stage from high stage tumors, whereas the trend was the same in the case of the Yoon, *et al.* data, it is likely that the n-value was too low to achieve significance (S3 Fig).

**Fig 1.**
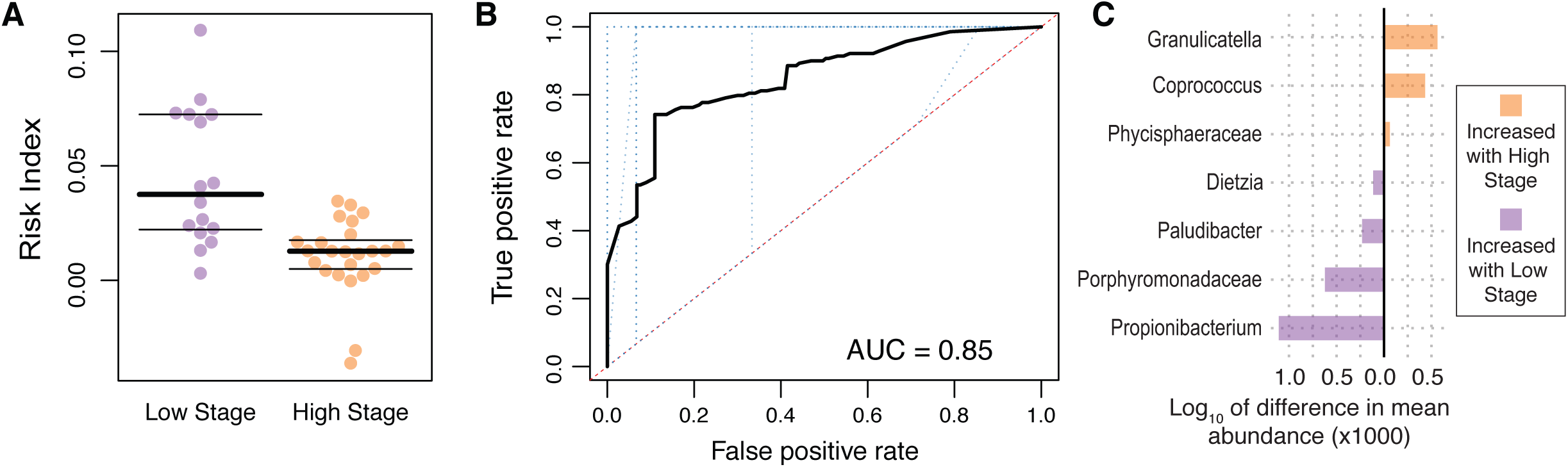
Correlation between the microbial community at a tumor that differentiates between tumor stage. (A) Low-stage (stages 1-2) and high-stage (stages 3-4) tumors can be differentiated using a risk index classifier generated from microbial abundance data (y-axis). The central black bar indicates the median, and the thin black bars represent the 25th and 75th percentiles. (B) A receiver operating characteristic (ROC) curve was generated using a 10-fold cross-validation (blue dotted lines). The average of the 10-fold cross-validation curves is represented as a thick black line. (C) Differences in the mean abundances of a subset of the taxa predicted to interact differentially with high-stage and low-stage tumors. This subset represents those taxa that had a mean difference in abundance of greater than 0.1%, proportionally.

The comparisons of interest here are between tumor samples at high stage and tumor samples at low stage. This study utilized patient-matched normal samples, but these are not true normals, as they came from the same cancer patients who themselves have low or high stage cancer. To address the question of how the CRC samples, grouped by stage, compare to independent normal samples from healthy individuals, we obtained unpublished colonic mucosal microbiome data from a separate, but otherwise methodologically similar, study being performed at the University of Minnesota that included the tissue-associated microbiome from individuals without cancer (n=12) undergoing routine colonoscopy. The OTU tables from the normal samples as well as the low-stage and high-stage samples were merged and assessed using the aforementioned LEfSe LOO approach (see Methods) to generate a model that is relevant to the normal and cancer samples (S4 Fig). The high-stage and low-stage samples were still able to be separated, as were the differences between the normal and the low-stage. Interestingly, the high-stage samples were not able to be statistically separated from the normal samples using this model.

### Tumor mutations correlate with consistent changes in the proximal microbiome

Next, we attempted to use a similar approach to classify tumors based on mutational profiles. We initially focused on individual genes that harbor loss-of-function (LoF) mutations, as those, we predicted, would be the most likely to have a physiologically relevant interaction with the surrounding microbiome. We applied a prevalence filter to include only those mutations that were present in at least 10 or more patients at the gene level, including 11 genes in the analysis. The purpose of the filter was to limit the number of statistical tests while maintaining a reasonable number of possibly cancer-driving mutations; see S7 Table for how changes in this cutoff affect the number of genes included in the analysis. The raw OTU table was collapsed to the level of genus for the analysis. A visualization of the correlations between gene-level mutational status and the associated microbial abundances revealed differing patterns of abundances that suggests an interaction between the 11 most prevalent LoF tumor mutations and the microbiome (**Fig. 2** and S5 Fig). We hypothesized that the presence of mutation-specific patterns of microbial abundances could be statistically described by prediction of tumor LoF mutations in individual genes using the microbiome. For each of eleven genes that passed prevalence filtering cutoff, we identified the associated microbial taxa (**Fig. 2A**, S6 Table, and S8 Table), generated risk indices for each patient (**Fig. 2B-C**), and plotted the mean differences in abundances for a subset of microbial taxa interacting with each mutation (**Fig. 2D**). We found that we are able to use microbiome composition profiles to predict the existence of tumor LoF mutations in the human genes *APC, ANKRD36C, CTBP2, KMT2C*, and *ZNF717* (Q-value = 0.0011, 0.0011, 0.019, 0.019, and 0.055, respectively, by permutation test after False Discovery Rate (FDR) correction for multiple tests; **Fig. 2**). The risk indices for each mutation were generated using sets of microbial taxa that ranges from 22 (*ZNF717*) to 53 (*ANKRD36C*) taxa (S6 Table). The taxa that showed the most dramatic differences in abundance when comparing tumors with and without mutations are shown in Fig 2D. For example, the abundance of *Christensenellaceae* is relatively lower in tumors with *APC* mutations, but relatively higher in tumors with *ZNF717* mutations.

**Fig 2.**
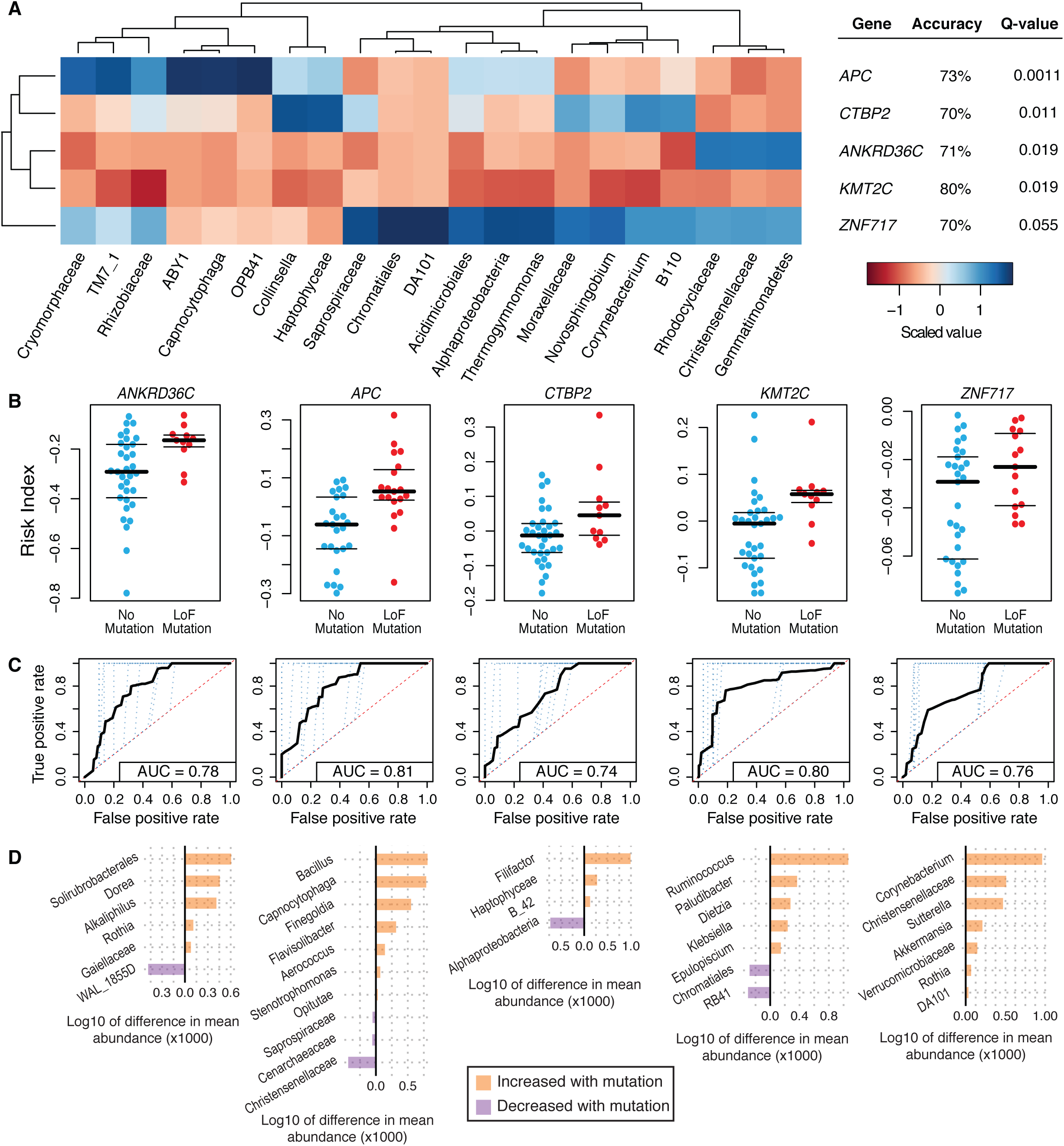
Commonly mutated genes show a predicted interaction with changes in the abundances of several microbial taxa. (A) A heatmap of the scaled abundances values (cells) for a subset of taxa chosen as they were identified as discriminatory in each leave-one-out iteration (columns) that were found significantly associated with prevalent LoF mutations (rows). Scaled abundances are from the patients with the indicated mutations. (B) LoF mutations in each of the indicated genes can be predicted using a risk index as a classifier (y-axis). The central black bar indicates the median, and the thin black bars represent the 25th and 75th percentiles. (C) ROC curves were generated for each of the indicated mutations using a 10-fold cross-validation (blue dotted lines). The average of the 10-fold cross-validation curves is represented as a thick black line. (D) Differences in the mean abundances of a subset of the taxa predicted to interact differentially with tumors with a LoF mutation relative to those without the indicated mutation. This subset represents those taxa that had a mean difference in abundance of greater than 0.1%, proportionally.

To determine if our ability to find associations between specific mutations and microbial signatures was simply the result of being confounded by stage (*e.g.* perhaps high-stage tumors all have mutations in *APC*, meaning that the ability to separate tumors by mutations in this gene are potentially just a function of stage), we used mutation presence/absence information and tumor stage to look for confounding correlations between the two using Fisher’s exact test. The results show that there is no confounding associations between mutation status and tumor stage (S9 Table).

Next, we applied our interaction prediction approach, as described above, to the pathway-level mutational data (**Fig. 3**; see Methods). Following visualization of the pathway level abundances (S6 Fig, S7 Fig), we found that LoF mutations in each of the 21 KEGG pathways passing prevalence filter can be significantly predicted with a fixed specificity of 80% and an accuracy up to 86% (Q-values < 0.02 by permutation test after FDR correction; **Fig 3A-D**, S10 Table, S8 Fig, and S9 Fig). Similarly, microbiome composition significantly predicted LoF mutations in 15 of the 19 tested PID pathways (Q-values < 0.04 by permutation test after FDR correction) (**Fig 3E-H,** S10 Table, S10 Fig, and S11 Fig). The full sets of taxon abundances that were specifically associated with each of the LoF mutations in the genes and pathways can be found in S11 Table, S12 Table, S13 Table, S14 Table, and S12 Fig. In general, the number of taxa within each of the sets used to generate the risk indices was lower than that used for the gene-level analyses, with an average of 37 taxa per gene compared to 7 taxa per pathway. When comparing results using the gene-level interactions and the pathway level interactions, for instance looking at mutations in *APC* (**Fig. 2**) and comparing them to mutations in the KEGG-defined Wnt signaling pathway and the PID-defined Canonical Wnt signaling pathway (**Fig 3**), the interactions at the pathway level are more statistically significant (AUC for *APC* = 0.81, KEGG = 0.88, PID = 0.90). This trend is consistent and can be visualized as a density histogram of interaction prediction accuracies (S13 Fig), indicating a stronger signal of interaction with the microbiome when using pathway-level mutational data.

**Fig 3.**
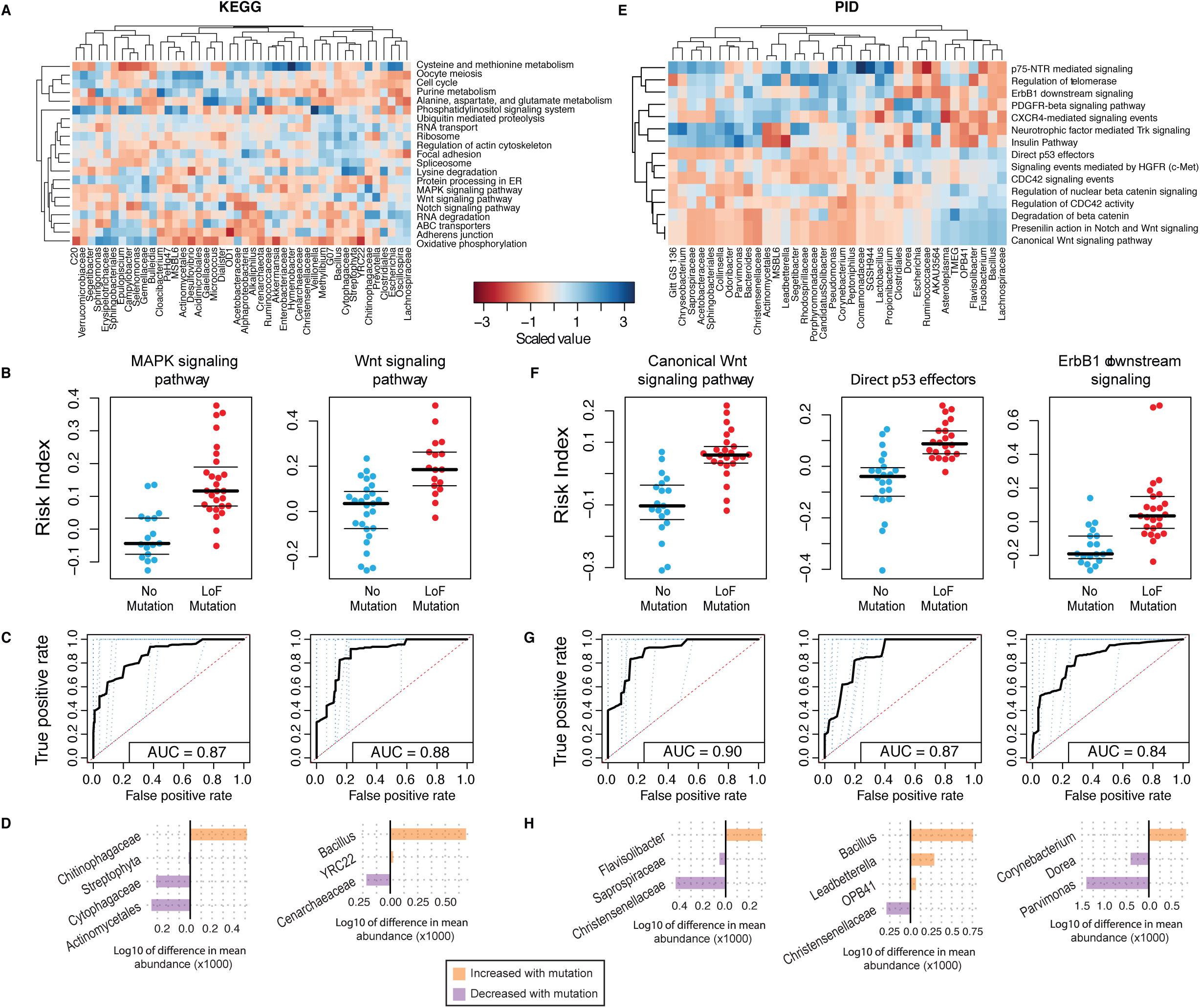
Pathways harboring prevalent LoF mutations correlate with changes in the abundances of sets of microbial taxa. (A) A heatmap of the scaled abundances values (cells) for a subset of taxa (columns) that are found significantly associated with KEGG pathways harboring LoF mutations (rows). Scaled abundances are from the patients with mutations in the indicated pathways. (B) LoF mutations in each of the indicated pathways can be predicted using a risk index as a classifier (y-axis). The central black bar indicates the median, and the thin black bars represent the 2nd and 4th quartiles. (C) ROC curves were generated for each of the indicated pathways using a 10-fold cross-validation (blue dotted lines). The average of the 10-fold cross-validation curves is represented as a thick black line. (D) Differences in the mean abundances of a subset of the taxa predicted to interact differentially with tumors harboring mutations in the indicated pathways relative to those without a mutation. This subset represents those taxa that had a mean difference in abundance of greater than 0.1%, proportionally. (E - F) Identically structured visualizations as in (A - D), but for PID pathway data rather than the KEGG pathways.

### Predicted microbiome interaction network affected by tumor mutational profile

Lastly, we assessed the correlations between taxa among tumors with and without LoF mutations (**Fig. 4**; see methods). We found striking differences in the structure of the network comparing tumors with and without a Lof mutation in *APC* the correlations between taxa (**Fig 4A**). For example, in tumors with mutations in *APC*, the abundance of *Christensenellaceae* is positively correlated with *Rhodocyclaceae* and negatively correlated with *Pedobacter*. In tumors lacking LoF mutations in *APC*, these correlations are lost and *Christensenellaceae* is instead negatively correlated with *Saprospiraceae* and *Gemm 1*. We also assessed the network of correlations across tumors with mutations in PID pathways (**Fig 4B**). This analysis highlighted that some pathway-level mutations show a shared set of correlations between taxa, while others appear independent. It is interesting to note that there are several PID pathways that biologically linked (*e.g.* Degradation of beta-catenin, Regulation of nuclear b-catenin signaling, Presenilin action in Notch and Wnt Signaling, and Canonical Wnt signaling pathway). These pathways do indeed share core elements, but as defined by the PID authors, are comprised of distinct combinations of genes (S14 Fig)[34]. Several of the taxa that can be used to predict LoF mutations in p75(NTR) signaling share correlations among each other as well as with taxa associated with mutations in PDGFR-beta signaling and direct p53 effectors.

**Fig 4.**
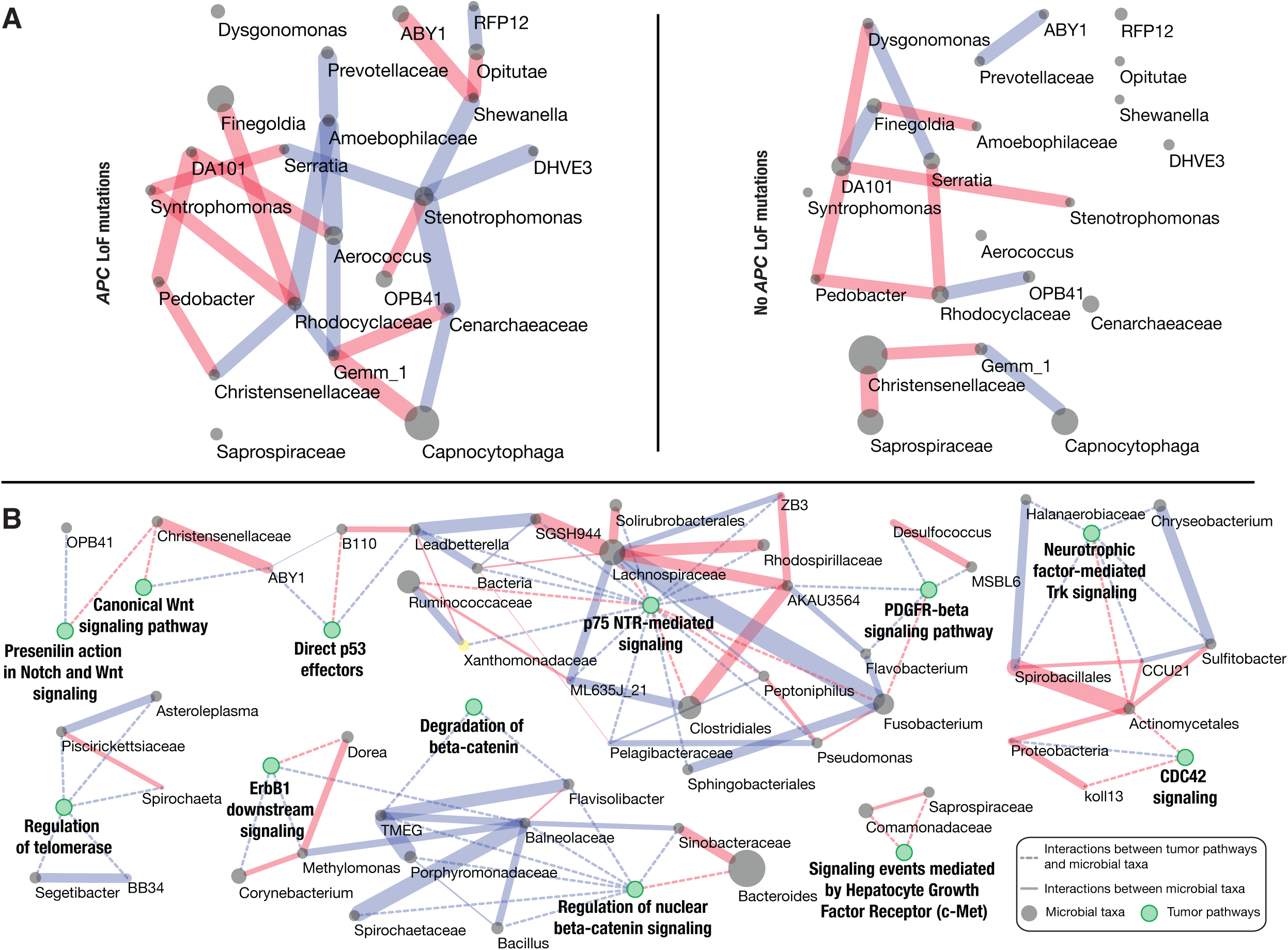
Interaction networks among bacteria are defined by host tumor mutations. (A) SparCC analysis of the microbial abundances of the taxa identified by LEfSe for tumors APCLoF mutations in APC (left) and without mutation (right) produce distinct patterns of correlations (edges) between a common set of taxa (nodes). Direct correlations are indicated as red edges and inverse correlations as blue edges (SparCC R >= 0.25, P <= 0.05 for displayed edges). (B) SparCC analysis was run simultaneously for all taxa identified by LEfSe when predicting interactions with mutations in PID pathways. There are interactions (dashed edges) between the taxa (grey nodes) associated with mutations across sets of PID pathways (green nodes). The solid edges indicate SparCC R-values (red for direct and blue for inverse correlations). The grey taxon nodes are scaled to the average abundance of the taxa in the associated tumor set. Edge color indicates the direction of the interaction, red for negative and blue for positive. Note that while several of the pathways (green nodes) have closely related general functions (*e.g.* “Canonical Wnt signaling pathway” and “Degradation of beta-catenin”), the underlying gene sets that comprise these pathways are distinct and result in independent correlations with microbial taxa.

## Discussion

The link between colorectal cancer and the gut microbiome has been highlighted by a large number of recent studies[2–17,19], with several hypotheses as to the causal role of microbes in the disease[9,12,43,44]. Given that host genetics is associated with microbiome composition, and since cancer is a genetic disease caused by mutations in host DNA, it is of interest to study the microbiome in the context of tumor mutational profiles[25–30]. Here, we jointly analyzed tumor coding mutational profile and the taxonomic composition of the proximal microbiome. We found that the composition of the microbiome is correlated with mutations in tumor DNA, and that this correlation can be used to statistically predict mutated genes and pathways solely based on the microbiome.

The association of microbial taxa with tumor stage (Fig 1) mirrors recent results, including a study of a Chinese population[4,41]. This concordance is relevant as it indicates that the microbial communities appear to be consistent even when comparing geographically distinct patient cohorts[45,46]. One of the predictive taxa, *Porphyromonadaceae*, has been shown to be altered in mouse models of CRC in other studies as well[7,14]. A study on the link between dysbiosis and colitis-induced colorectal cancer also showed similar results[47]. For instance, the bacterial genus *Paludibacter* was found to be associated with risk of developing tumors in a mouse model[47]. Additionally, other researchers have identified this taxa as a possible contributor to inflammation in bovine mastitis[48]. We find that *Paludibacter* is significantly associated with low-stage tumors, again, supporting the hypothesis that these bacteria are associated with cancer risk and may be contributing to early stage inflammation[47,48]. An equally likely hypothesis is that *Paludibacter* is using the inflammatory microenvironment to its own advantage, thereby explaining its association with the tissues at this state. Conversely, we found that the genus *Coprococcus* is associated with high-stage tumors and not low stage tumors. Members of this genus are known to generate butyrate and propionate, short chain fatty acids (SCFAs) that can be utilized by host colonocytes as an energy source and which, in this context, can act as anti-inflammatory SCFAs[49,50]. Although our results are correlational and cannot point to causal effects, these findings suggest that driving inflammation may play a role in early stage cancer, while generating nutrients at the cost of suppressing inflammation may be more beneficial to the tumor in later stages. As microbes can have different effects dependent on their environmental contexts, further work that studies the overall functional effects of microbial communities will need to be performed to assess how individual members interact and contribute to, suppress, or have no influence on inflammation.

Gene-level mutation data, visualized in S5 Fig, show intriguing patterns of microbial abundances that are associated with the tumors harboring different mutations. While our previous work demonstrated that specific taxa, including *Providencia*, were significantly associated with tumors when compared to patient-matched normals, we did not find that this genus significantly discriminated among tumors based on LoF mutation status[2]. This suggests that there are potentially microbial taxa that might act as generic tumor-associated microbes and others that might rely on specific alterations in the tumor. For instance, as reflected in the differing patterns within each gene (rows) in the heatmap, *Aerococcus* and *Dorea* both show higher abundances within tumors harboring LoF mutations in *ZNF717, CTBP2*, and *APC*, relative to tumors with LoF mutations in *ANKRD36C* and *KMT2C*. This highlights the different patterns in the microbiome that can be found when assessing genetically heterogeneous sets of tumors, as *Dorea* has been found to be differentially present in tumor microbiomes by several different groups. Our work highlights some potential genetic interactions that may explain the differences seen (*e.g.* differences in the mutational statuses of the patients in the different cohorts could result in different findings)[3,5–8]. Thus, incorporating host genetic profiles in studies of the microbiome in CRC may be beneficial and uncover patterns in clearly specified subsets of patients that are defined depending on specific tumor mutations.

While it is not possible to definitively identify the biological mechanism behind the predicted interactions among mutated genes and microbial taxa due to the correlative nature of this work (shown in Fig 2), it is possible to generate hypotheses based on what is already known in the relevant literature. For example, we found that LoF mutations in *APC* correlate with changes in 25 different microbial taxa, including an increase in the abundance of the genus *Finegoldia*. This genus was identified in previous studies of colon adenomas and harbors species that are opportunistic pathogens at sites of epithelial damage[6,51,52]. *Capnocytophaga* has been identified as a potential biomarker for lung cancer[53]. Our results also indicate that changes in the abundance of *Christensenellaceae* are associated with mutations in both *APC* and *ZNF717*. A recent study in twins has identified *Christensenellaceae* as a taxon that is highly driven by host genetics[27]. We found that mutations in *ZNF717*, a transcription factor commonly altered in gastric, hepatocellular, and cervical cancers[54–56], are associated with *Verrucomicrobiaceae* and *Akkermansia*, which are both known to increase in abundance in conjunction with colitis[57]. *Alphaproteobacteria* are significant contributors to our ability to predict mutations in *CTBP2*, a repressor of transcription known to interact with the ARF tumor suppressor[58]. Changes in this bacterial taxon’s abundance has also been found to be associated with prostate cancer, although the mechanism of action is unknown[59]. We also show that mutations in *KMT2C*, a gene commonly co-mutated along with *KRAS,* could be predicted, in part, using the abundance of *Ruminococcus[60]*. These bacteria have been previously implicated in inflammatory bowel disorders and colorectal cancer by multiple groups[8,61–63].

Similar results were also evident when aggregating the mutations into KEGG and PID pathways (Fig 3, S6 Fig, and S7 Fig; see Methods)[31–34]. As an example, we found that the abundance of microbes that predict mutations in KEGG pathways form two distinct clusters, and that the genus *Escherichia* has a higher scaled abundance in tumors with mutations in the KEGG pathways in cluster 1 relative to those in cluster 2 (S6 Fig). Cluster 1 contains adherens junctions, which are partially responsible for maintaining the intestinal barrier; a disruption of the intestinal barrier in mice using cyclophosphamide was shown to cause a loss of adherens junction function and a concomitant increase in bacterial translocation into the intestinal tissue, including species of *Escherichia[64]*. When examining the heatmap with LoF mutation collapsed into PID pathways (S7 Fig), we again identified differences in scaled microbial abundances between the tumors as a function of which pathways are mutated. For instance, we found lower abundance of *Pseudomonas* in tumors with LoF mutations in the pathways ‘regulation of nuclear β-catenin signaling and target gene transcription’, ‘degradation of β-catenin’, ‘presenilin action in Notch and Wnt signaling’, and ‘canonical Wnt signaling pathways’. Recent studies have shown that *Pseudomonas* strains that express the *LecB* gene can lead to degradation of β-catenin, providing hypothetical support for the concept that this genus may play a somewhat protective role in CRC by suppressing the Wnt signaling pathway[65]. The mechanism that might explain this phenomenon is still unclear, but may have to do with alterations in appropriate cell surface adhesion molecules for the LecB protein or a change in the content of the cellular microenvironment[65,66].

Many of the interactions identified here between bacterial taxa and mutations in PID pathways have already been demonstrated experimentally in the literature. For example, in human oral cancer cells, it was shown that bacteria of interest were able to activate EGFR through the generation of hydrogen peroxide[67]. In addition, the correlation between ErbB1 downstream signaling and increase in the abundance of *Corynebacterium* has been demonstrated mechanistically in a model of atopic dermatitis, whereby EGFR inhibition results in dysbiosis (the appearance of *Corynebacterium* species) and inflammation[68]. Specific depletion of *Corynebacterium* ablates the inflammatory response[68]. Moreover, our finding that the abundance of *Fusobacterium* is depleted in tumors with LoF mutations in the PDGFR-beta pathway may be explained by the dependence of several pathogenic strains of bacteria for functionally intact PDGFR signaling for adherence to intestinal epithelium[69]. In addition, p75(NTR) signaling has been shown to operate as a tumor suppressor by mediating apoptosis in response to hypoxic conditions and reactive oxygen species[70–73]. Alterations in this pathway have also been shown to be useful as a biomarker for esophageal cancer[74,75].

Our study has several caveats. First, our study only shows correlations, and we cannot directly assess causal effects. Thus, we do not know whether the microbiome is altered before or after the appearance of specific mutations. Nevertheless, many of the predicted interactions described above have been previously tested, albeit across a wide variety of experimental systems and disease states, typically in isolation, for biological relevance and mechanism of action. We anticipate that future studies will comprehensively test the causality of interactions by utilizing model organisms and cell culture techniques, where the directionality of the interactions between mutations and microbial taxa can be assessed. Additionally, we have only profiled the taxonomic composition of the microbiome, and thus cannot detect interactions that are dependent on microbial genes or functions. We also do not have an exhaustive history of each of the patients and their backgrounds (*e.g.* diet, family history, immune profile, *etc*.), without which is it possible there may be a confounding variable that explains some of our correlations. Again, this work is the requisite precursor to functional analyses in a wet-lab setting. Similarly, using whole-exome sequencing does not allow us to include non-coding mutations and larger tumor structural variants and chromosomal abnormalities. This can be alleviated by the use of metagenomic shotgun sequencing to profile the microbiome, as well as whole-genome sequencing to assess tumor mutations. Moreover, the study sample was relatively small (n = 88 samples from 44 patients). Nevertheless, the sample size was sufficient to detect significant patterns. As ours is the first study, as far as we know, to present simultaneous exome sequencing of tumors alongside assessment of the microbial communities associated with these tumors, no validation datasets are available. Additional studies that use large tumor samples would be useful in validating our results and identifying further associations as targets of functional interaction assays.

From a broad perspective, tumors arise and thrive as a result of the combinatorial influences of their own genetics and their environments. In the context of the microbiome, there are three potential roles that microbial taxa may be playing in tumor biology: (1) they can contribute to the development and/or progression of the tumor; (2) they might have no effect on tumor formation or development, acting as pure commensals; and (3) they may provide protection against tumor development or progression. The research presented here does not address which of these hypotheses are most likely to play the largest role, as it is highly probable that there are examples of microbial communities that align with each of these examples. The work presented here is a first attempt at identifying which points in the interplay between tumor genetics and the microbes in the tumor environment are ripe targets for future analysis to ascertain function and definitively identify what role these taxa are playing in the context of cancer pathogenesis, if any.

In summary, we present an association between tumor genetic profiles and the proximal microbiome, and identify tumor genes and pathways that correlate with specific microbial taxa. We also show that the microbiome can be used as a predictor of mutated genes and pathways within a tumor, and suggest potential mechanisms driving the interaction between the tumor and its microbiota. Our proof-of-principle analysis can provide a starting point for the development of diagnostics that utilize microbiome profiles to ascertain CRC tumor mutational profiles, facilitating personalized treatments.

## Materials and Methods

### Patient inclusion and DNA extraction

88 tissue samples from 44 individuals were used, with one tumor and one normal sample from each individual. These de-identified samples were obtained from the University of Minnesota Biological Materials Procurement Network (Bionet), a facility that archives research samples from patients who have provided written, informed consent. These samples were previously utilized and are described in detail in a previous study[76]. The patient information provided for this retrospective cohort did not include information related to immune status or inflammation. To reiterate these points, all research conformed to the Helsinki Declaration and was approved by the University of Minnesota Institutional Review Board, protocol 1310E44403. Tissue pairs were resected concurrently, rinsed with sterile water, flash frozen in liquid nitrogen, and characterized by staff pathologists. The criteria for selection were limited to the availability of patient-matched normal and tumor tissue specimens. In all cases, normal tissue was confirmed by the pathologist to be tumor free. The tumor stages were classified using a modified stage grouping based on the TNM scale (collapsing the letter-scales into single numbers - eg. IIIA, IIIB, and IIIC are reported as stage 3. Additional patient metadata are provided in S4 Table and in the indicated work[76]. Microbial community data from 16S rRNA gene sequencing of an independent cohort of healthy patients (n =12) who underwent routine colonoscopies was acquired from a separate, unpublished dataset and used in the comparison of low-stage, high-stage, and normal microbial communities.

### Microbiome characterization

The microbiome data used in the study was generated previously and is described exhaustively in[76]. Briefly, microbial DNA was extracted from patient-matched normal and tumor tissue samples using sonication for lysis and the AllPrep nucleic acid extraction kit (Qiagen, Valencia, CA). The V5-V6 regions of the 16S rRNA gene were PCR amplified with the addition of barcodes for multiplexing using the forward and reverse primer sets V5F and V6R from Cai, et al.[77]. The barcoded amplicons were pooled and Illumina adapters were ligated to the reads. A single lane on an Illumina MiSeq instrument was used (250 cycles, paired-end) to generate 16S rRNA gene sequences. The sequencing resulted in approximately 10.7 million total reads passing quality filtering in total, with a mean value of 121,470 quality reads per sample. The forward and reverse read pairs were merged using the USEARCH v7 program ‘fastq_mergepairs’, allowing stagger, with no mismatches allowed[78]. OTUs were picked using the closed-reference picking script in QIIME v1.7.0 using the Greengenes database (August 2013 release)[79–81]. The similarity threshold was set at 97%, reverse-read matching was enabled, and reference-based chimera calling was disabled.

### Exome sequence data generation

Genomic DNA samples were quantified using a fluorometric assay, the Quant-iT PicoGreen dsDNA Assay Kit (Life Technologies, Grand Island, NY). Samples were considered passing quality control (QC) if they contained greater than 300 nanograms (ng) of DNA and display an A260:280 ratio above 1.7. Full workflow details for library preparation are outlined in the Nextera Rapid Capture Enrichment Protocol Guide (Illumina, Inc., San Diego, CA). In brief, libraries for Illumina next-generation sequencing were generated using Nextera library creation reagents (Illumina, Inc., San Diego, CA). A total of 50 ng of genomic DNA per sample were used as input for the library preparation. The DNA was tagmented (simultaneously tagged and fragmented) using Nextera transposome based fragmentation and transposition as part of the Nextera Rapid Capture Enrichment kit (Illumina, Inc., San Diego, CA). This process added Nextera adapters with complementarity to PCR primers containing sequences that allow addition of Illumina flow cell adapters and dual-indexed barcodes. The tagmented DNA was amplified using dual indexed barcoded primers. The amplified and indexed samples were pooled (8 samples per pool) and quantified to ensure appropriate DNA concentrations and fragment sizes using the fluorometric PicoGreen assay and the Bioanalyzer High-Sensitivity DNA Chip (Agilent Technologies, Santa Clara, CA). Libraries were considered to pass QC as long as they contained more than 500 ng of DNA and had an average peak size between 200 - 1000 base pairs. For hybridization and sequence capture, 500 nanograms of amplified library was hybridized to biotinylated oligonucleotide probes complementary to regions of interest at 58° C for 24 hours. Library-probe hybrids were captured using streptavidin-coated magnetic beads and subjected to multiple washing steps to remove non-specifically bound material. The washed and eluted library was subjected to a second hybridization and capture to further enrich target sequences. The captured material was then amplified using 12 cycles of PCR. The captured, amplified libraries underwent QC using a Bioanalyzer, and fluorometric PicoGreen assay. Libraries were considered to pass QC as long as they contained a DNA concentration greater than 10 nM and had an average size between 300 - 400 base pairs. Libraries were hybridized to a paired end flow cell at a concentration of 10 pM and individual fragments were clonally amplified by bridge amplification on the Illumina cBot (Illumina, Inc., San Diego, CA). Eleven lanes on an Illumina HiSeq 2000 (Illumina, Inc., San Diego, CA) were required to generate the desired sequences. Once clustering was complete, the flow cell was loaded on the HiSeq 2000 and sequenced using Illumina’s SBS chemistry at 100 bp per read. Upon completion of read 1, base pair index reads were performed to uniquely identify clustered libraries. Finally, the library fragments were resynthesized in the reverse direction and sequenced from the opposite end of the read 1 fragment, thus producing the paired end read 2. Full workflow details are outlined in Illumina’s cBot User Guide and HiSeq 2000 User Guides. Base call (.bcl) files for each cycle of sequencing were generated by Illumina Real Time Analysis (RTA) software. The base call files and run folders were then exported to servers maintained at the Minnesota Supercomputing Institute. Primary analysis and de-multiplexing was performed using Illumina’s CASAVA software 1.8.2. The end result of the CASAVA workflow was de-multiplexed FASTQ files that were utilized in subsequent analysis for read QC, mapping, and mutation calling.

### Exome data analysis

The exome sequence data contained approximately 4.2 billion reads in total following adapter removal and quality filtering, inclusive of forward and reverse reads, with a mean value of 47.8 million high-quality reads per sample. The raw reads were assessed using FastQC v0.11.2 and the Nextera adapters removed using cutadapt v1.8.1[82,83]. Simultaneously, cutadapt was used to trim reads at bases with quality scores less than 20. Reads shorter than 40 bases were excluded. The trimmed and filtered read pairs were aligned and mapped to the human reference genome (hg19) using bwa v0.7.10 resulting in a bam file for each patient sample[84]. These files were further processed to sort the reads, add read groups, correct the mate-pair information, and mark and remove PCR duplicates using picard tools v1.133 and samtools v0.1.18[85,86]. Tumor-specific mutations were identified using FreeBayes v0.9.14-24-gc292036[87]. Following these steps, 94.0% of the remaining read pairs mapped to the reference genome, hg19. Specifically, SNPs-only were assessed and a minimum coverage at each identified mutation position of more than 30X was required in both the patient normal and tumor samples. These mutations were filtered to only include those that were within protein-coding regions which were then compiled into a single vcf file. We did not remove variants found in dbSNP. This vcf file was assessed using SNPeffect v4.1 K (2015-09-0) in order to predict the potential impact of each of the mutations[88]. Based on these results, the mutations were grouped into three categories: (1) total mutations (2) non-synonymous mutations and (3) loss of function (LoF) mutations. The total mutations group The total mutations group is simply the sum of all the mutations that were found in each tumor. The non-synonymous mutations included all the mutations in the total mutations group that were non-silent. The LoF group only included those mutations that resulted in a premature stop codon, a loss of a stop codon, or a frameshift mutation.

Mutations in genes were collapsed to pathways (PID and KEGG) based on pathway membership as defined by the relevant database authors[31,33,34]. Specifically, we relied upon Uniprot annotations to link genes to pathways with the database file, idmapping.dat.gz, available from Uniprot[89,90]. This, three column database file is used by the web-based mapping interface for cross-referencing between genes and pathway membership datasets, including KEGG and PID[89]. In our case, we annotated each gene in our dataset with pathway membership information by first reducing the size of the idmapping.dat file by including only entries for KEGG and PID. For each LoF mutation in our dataset, we found the corresponding gene in the reduced database file with a matching ENSEMBL Gene ID (S1 Table) and recorded which grouping it was found in (KEGG or PID) and what the name of the defined pathway was (*e.g.* Canonical Wnt signaling). Many genes had multiple annotations as they are members of more than one of the defined pathways. These annotations were used to identify which tumor samples had mutations in KEGG and PID pathways, defined as having at least one gene harboring a LoF mutation that was a member of the pathway.

### Joint analysis of microbiome data with tumor stage and mutation status in genes and pathways

In order to identify microbial taxa that were significantly associated with specific characteristics (*i.e.* tumor stage - Fig 1, LoF mutations in individual genes - Fig 2, and LoF mutations within functional pathways - Fig 3), the OTU table was divided into two groups of tumors, defined by the characteristic of interest (*e.g.* Low Stage vs. High Stage, Mutation vs. No Mutation). As it would be invalid to generate a model that was built using the actual test sample, we used a “leave-one-out” approach when generating risk indices. To generate a risk index for a tumor, we started with an OTU table that contained all the tumors (43), leaving out the one for which we were calculating the risk index. This 43-tumor OTU table was divided into two groups based on the characteristic of interest, as described above. These data were used as input into LEfSe, as a means of identifying the microbial taxa that were able to discriminate between the two groupings[40]. Taxa were considered significant discriminators if the base 10 logarithm of their LDA score was less than 2, as recommended. The taxa that were significant discriminators were themselves of two categories: those that were more abundant, for instance, at tumors that harbored a mutation and those that were more abundant at tumors that did not harbor a mutation. The relative abundances of these taxa were arcsine root transformed and these transformed values - one for each category - were summed. The difference between these sums was calculated and is the risk index value used in each of the aforementioned figures. The use of the unweighted sum in the risk index, rather than relying entirely on the regression coefficients from LDA, is a simple way to control the degree of flexibility of the model when training on small sample sizes. More detail is described in a previous publication[91]. This procedure was repeated a total of 44 times (once for each tumor, to complete the leave-one-out approach) to obtain a risk index for each of the patients. The significance of the difference in risk indices between the patients found in one group vs. another (*e.g.* low stage vs. high stage, with LoF mutation, without LoF mutation) was assessed using a Mann-Whitney U test and a permutation test, in which we permuted the labels for a given group 999 times, each time deriving new held-out predictions of the risk indexes for each subject for that gene. Then the observed difference in means between the patients with LoF mutation and patients with LoF mutation risk index predictions using the method on the actual LoF mutation labels to the differences observed in the permutations to obtain an empirical P-value was compared. The resulting P-values were corrected using the false discovery rate (FDR) correction for multiple hypothesis tests.

Receiving Operating Characteristic (ROC) curves were plotted and the area under the curve (AUC) values computed on a dataset containing 10 sets of predictions and corresponding labels obtained from 10-fold cross-validation using ROCR package in R[92]. A risk index threshold was also obtained that best predicts the membership in a stage group or the presence or absence of LoF mutation with a leave-one-out cross-validation on the risk index. Each held-out sample was treated as a new patient on whom the optimal risk index cutoff was tested and subsequently refined to separate patients who had a LoF mutation and patient who did not have a LoF.

Correlation analysis was performed using SparCC on a reduced OTU table containing significant taxa identified using the above prediction methods collapsed to the genus level[93]. Pseudo p-values were calculated using 100 randomized sets. Networks of correlations were visualized using Cytoscape v3.1.0[94].

The patients in this study have associated clinical data as we described previously[76], We used a linear model to determine the extent to which clinical factors may correlate with mutation load. These included patient sex, tumor stage, patient age, patient body mass index (BMI), and microsatellite instability (MSI) status. None of these factors, alone or in combination, were found to significantly impact the mutational data, though it bears noting that MSI status was only available for a subset (13 out of 44) of the patients.

### Availability of data and materials

The microbiome datasets supporting the conclusions of this article are available in the NCBI sequence read archive: project accession number PRJNA284355, https://www.ncbi.nlm.nih.gov/sra/PRJNA284355.

The Exome sequencing dataset supporting the conclusions of this article are available (with proper access approval) from dbGAP - [*project accession number and hyperlink currently being generated*].

The tumor-specific mutation data supporting the conclusions of this article is included as S1 Table.

## Acknowledgments

We thank the members of the Blekhman Lab for helpful discussions. This work was supported, in part, by resources provided by the Minnesota Supercomputing Institute.

## Author Contributions

M.B.B. and R.B. designed the project. M.B.B. performed DNA extractions, exome sequencing analysis, microbiome sequencing analysis, prepared the manuscript, and generated the figures. E.M. and M.B.B performed statistical analyses including LEfSe, the LOO approach, ROC curve generation, and permutation tests. J.A. assisted with the exome sequencing data analysis pipeline. S.P. performed initial data analysis of the independent normal microbiome sample set. D.E.N. and A.K. provided the independent normal samples with microbiome sequencing. A.K. contributed to the data interpretation and provided clinical guidance. T.K.S. contributed to the experimental design and interpretations of data. D.K. contributed to the design and interpretation of the interaction prediction approach. R.B. provided guidance related to the analytical approaches and interpretation of the results. All authors contributed to manuscript and figure revisions.

## Supplementary Information

**S1 Table**

Tumor specific mutations. This files is a large, tab-separated dataset that contains information on each of the tumor-specific mutations found in the patient exome sequences.

**S2-S16 Tables.**

An excel file contains S2-S16 Tables, each with their own descriptive headers, as individual tabs. This information includes specifics related to the exome sequencing, patient metadata, sets of taxa identified by LEfSe, and ROC curve generation.

**S1 Fig.**

Histogram of the proportion of reads passing quality filter that aligned to the references genome.

**S2 Fig.**

Tumors harbored a wide range of mutations. The histograms at top indicate the frequency distribution of the total (green), missense (blue), and LoF (red) mutations found in the tumors from our patient cohort. The plots at bottom show the absolute numbers of tumor-specific mutations detected in the tumors on a flat scale (bottom left) as well as on a log scale (bottom right).

**S3 Fig.**

Column dot plots of the risk indices determined using the stage-relevant model generated in this work to the microbiome data from Flemer, et al. (left) and Yoon, et al. (right).

**S4 Fig.**

Column dot plots of the risk indices calculated using a new model designed to separate microbial communities from normal, low stage, and high stage tumors. Normals included in this model and figure are from healthy individuals from an independent study, not from patient matched samples.

**S5 Fig.**

Heatmaps demonstrating the patterns of microbial abundances for patient samples with prevalent LoF mutations.

(A) Scaled taxon abundances (columns) in the tumor samples that harbor LoF mutations in the genes indicated (rows).

(B) Scaled differences (tumor abundance - matched normal abundance) patients that harbor tumor-specific LoF mutations in the genes indicated (rows).

**S6 Fig.**

Heatmaps demonstrating the patterns of microbial abundances for patient samples with prevalent LoF mutations in KEGG pathways.

(A) Scaled taxon abundances (columns) in the tumor samples that harbor LoF mutations in the KEGG pathways indicated (rows). Clusters 1 and 2 are labeled to facilitate discussion in main text.

(B) Scaled differences (tumor abundance - matched normal abundance) patients that harbor tumor-specific LoF mutations in the KEGG pathways indicated (rows).

**S7 Fig.**

Heatmaps demonstrating the patterns of microbial abundances for patient samples with prevalent LoF mutations in PID pathways.

(A) Scaled taxon abundances (columns) in the tumor samples that harbor LoF mutations in the PID pathways indicated (rows).

(B) Scaled differences (tumor abundance - matched normal abundance) patients that harbor tumor-specific LoF mutations in the PID pathways indicated (rows).

**S8 Fig.**

LoF mutations in KEGG pathways can be predicted using a risk index as a classifier (y-axis).

**S9 Fig.**

ROC curves were generated for each of the KEGG pathways indicated in S6 Fig using a 10-fold cross-validation (blue dotted lines). The average of the 10-fold cross-validation curves is represented as a thick black line. The AUC are indicated at bottom of each pathway panel.

**S10 Fig.**

LoF mutations in PID pathways can be predicted using a risk index as a classifier (y-axis).

**S11 Fig.**

ROC curves were generated for each of the PID pathways indicated in S8 Fig using a 10-fold cross-validation (blue dotted lines). The average of the 10-fold cross-validation curves is represented as a thick black line. The AUC are indicated at bottom of each pathway panel. This figure also includes BARD1 and Class I PI3K pathways, for references, although neither of these achieved statistical significance in other tests.

**S12 Fig.**

Large set of abundance plots for the taxa from each of the genes, KEGG pathways, and PID pathways that harbored prevalent LoF mutations and showed significance as a means of predicting the interaction between the mutation and the microbiota. Abundances are plotted as both column dot plots as well as horizontal bar plots of the differences in the mean abundances of a subset of the taxa predicted to interact differentially with tumors with a LoF mutation relative to those without the indicated mutation. These subsets represent those taxa that had a mean difference in abundance of greater than 0.1%, proportionally.

**S13 Fig.**

Interaction prediction accuracies increase when assessing biological pathways. A density histogram showing the distribution of prediction accuracies for individual genes (red), KEGG pathways (green), and PID pathways (blue). Interaction prediction accuracies are highest for human cancer-specific pathways from PID. The genes (red) exhibit a double peak distribution due to the relatively high accuracy achieved when predicting the presence of LoF mutations in *KMT2C* (80%).

**S14 Fig.**

Plot demonstrating the co-membership of genes within PID pathways. Horizontal bar chart indicates the total numbers of genes within each of the 4 PID pathways. Vertical bars in the bar chart indicate numbers of genes that fall within the memberships indicated by filled in dots and lines at the intersections below the bars.

